# Ligand Effects on Phase Separation of Multivalent Macromolecules

**DOI:** 10.1101/2020.08.15.252346

**Authors:** Kiersten M. Ruff, Furqan Dar, Rohit V. Pappu

## Abstract

Biomolecular condensates enable spatial and temporal control over cellular processes by concentrating biomolecules into non-stoichiometric assemblies. Many condensates form via reversible phase transitions of condensate-specific multivalent macromolecules known as scaffolds. Phase transitions of scaffolds can be regulated by changing the concentrations of ligands, which are defined as non-scaffold molecules that bind to specific sites on scaffolds. Here, we use theory and computation to uncover rules that underlie ligand-mediated control over scaffold phase behavior. We use the *stickers*-and-*spacers* model wherein reversible non-covalent crosslinks among stickers drive phase transitions of scaffolds, and spacers modulate the driving forces for phase transitions. We find that the modulatory effects of ligands are governed by: the valence of ligands; whether they bind directly to stickers versus spacers; and the relative affinities of ligand-scaffold versus scaffold-scaffold interactions. In general, all ligands have a diluting effect on the concentration of scaffolds within condensates. Whereas monovalent ligands destabilize condensates, multivalent ligands can stabilize condensates by binding directly to spacers or destabilize condensates by binding directly to stickers. Bipartite ligands that bind to stickers and spacers can alter the structural organization of scaffold molecules within condensates even when they have a null effect on condensate stability. Our work highlights the importance of measuring dilute phase concentrations of scaffolds as a function of ligand concentration in cells. This can reveal whether ligands modulate scaffold phase behavior by enabling or suppressing phase separation at endogeneous levels thereby regulating the formation and dissolution of condensates *in vivo*.

**Significance:** Phase transitions of multivalent macromolecules known as scaffolds help drive the formation of functional biomolecular condensates in cells. The formation and dissolution of condensates is tightly regulated, as aberrant phase behavior is associated with disease. Here, we show that distinct types of ligands can exert control over the formation and dissolution of condensates by binding to distinct sites on scaffold molecules. We further show that the extent and direction of regulation can be inferred through direct measurements of how ligands impact scaffold phase boundaries. Our findings have broad implications for understanding and modeling ligand-mediated regulation of condensates in cells, and for designing novel molecules that exert regulatory control over condensates.

---

Membraneless biomolecular condensates concentrate biomolecules in cells to organize biochemical reactions in space and time (1, 2). There is growing evidence that condensates form via spontaneous or driven phase transitions (2, 3). Functional condensates can be reconstituted *in vitro* and manipulated in live cells using only one or a small category of macromolecular scaffolds (4–13). Multivalence of interaction motifs known as *stickers* is a defining hallmark of scaffolds that drive phase transitions (8, 9) by combining density transitions in the form of phase separation and networking transitions in the form of percolation (4, 14, 15).

Mutations to scaffold molecules are associated with disease and evidence is growing that these lead to changes in scaffold phase behavior (16). Changes include lowering the threshold scaffold concentration needed for phase separation and lowering the barrier for liquid-to-solid transitions within condensates as summarized in **Table S1** of the *SI Appendix*. These results suggest that the formation and dissolution of condensates has to be tightly regulated in cells. One route to modulating phase behavior is through post-translational modifications to scaffold molecules (17). A second mechanism takes advantage of the fact that condensates contain several types of non-scaffold molecules. The expression levels of these non-scaffold molecules can be used as knobs that can be turned to control scaffold phase behavior (**Fig. 1**, **Table S2**, *SI Appendix*). We define these modulatory non-scaffold molecules as ligands.

**Figure 1:**
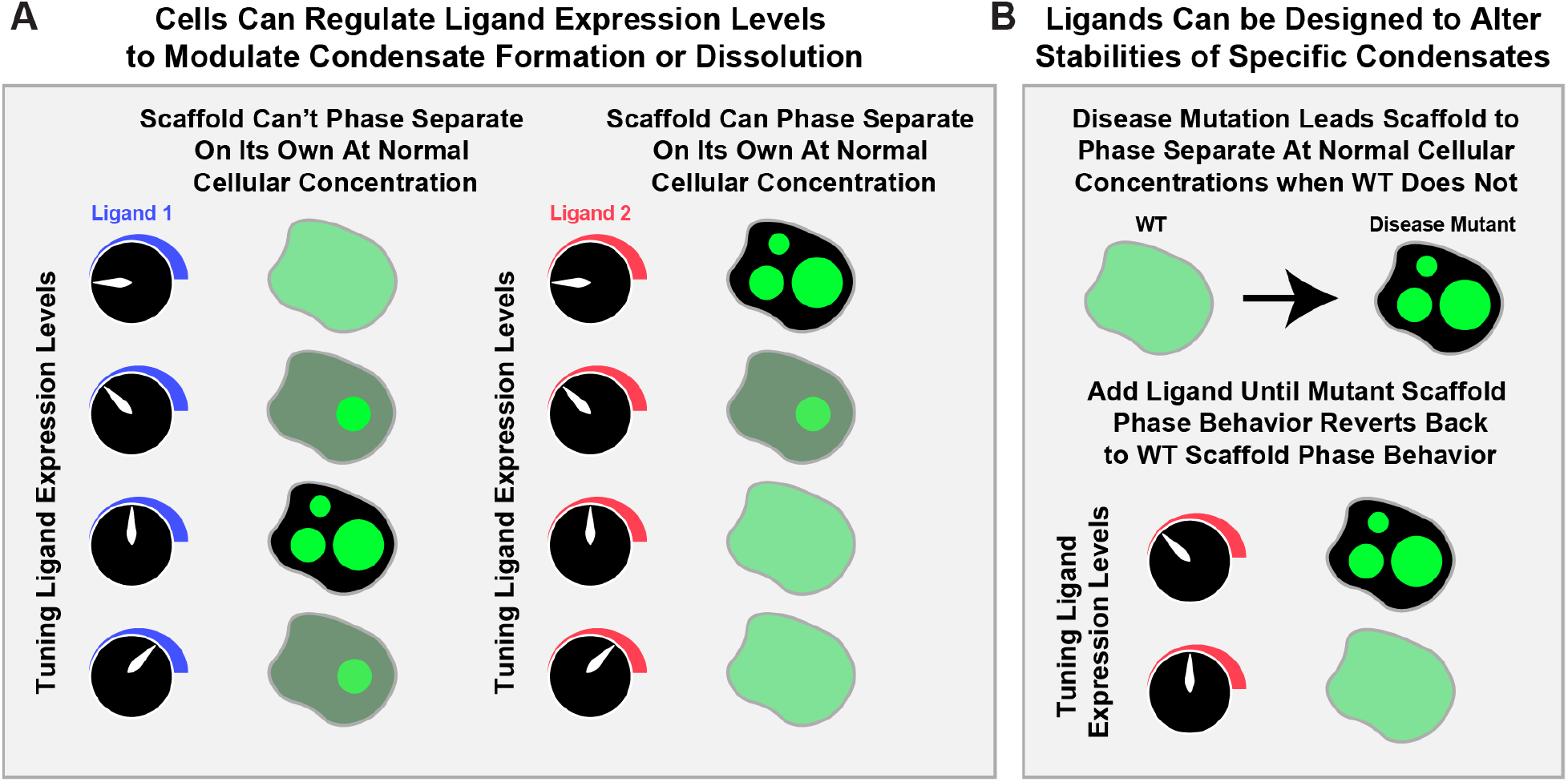
Schematic showing how the concentrations of preferentially binding ligands can modulate scaffold phase behavior in cells. Here, ligand expression levels are tuned from zero to high expression levels and green circles imply condensate formation of a fluorescently tagged scaffold. Hence, the brightness denotes the relative concentration of the scaffold in each phase.

Ligands do not undergo phase separation on their own and are not required for the phase separation of scaffolds. However, they bind preferentially to scaffolds across phase boundaries and either promote or destabilize condensates in cells. This is important because in some situations, the cellular concentrations of scaffolds may be too low to drive phase separation (18). This can be remedied by the controlled expression of specific ligands that lower the concentration threshold for phase separation (**Fig. 1A**). Alternatively, for scaffolds that can phase separate on their own at endogenous concentrations, increasing the expression level of a destabilizing ligand can help dissolve the condensate (19) (**Fig. 1A**). Similar ideas can be brought to bear in designing pharmaceutical approaches to regulate condensates (**Fig. 1B**).

To understand the mechanisms that underlie the regulation of scaffold phase behavior by ligands, we coopt the *polyphasic linkage* formalism of Wyman and Gill (20). Although this formalism was introduced four decades ago, it has not been deployed to understand, interpret, or appreciate the true scope of ligand-modulated phase separation *in vitro* and in live cells. Here, we establish why polyphasic linkage is useful for understanding how ligands can alter scaffold phase behavior. This, as explained above, is of direct relevance for understanding biological and pharmaceutical regulation of condensates *in vivo* (**Fig. 1**).

To illustrate the concepts of polyphasic linkage, we consider an aqueous solution with a single type of scaffold that separates into two distinct phases. We denote the scaffold-deficient phase as *A* and the coexisting scaffold-rich phase as *B*. The binodal or coexistence curve delineates the two-phase regime (**Fig. S1**, *SI Appendix*). For a given set of solution conditions, quantified in terms of an effective interaction strength, the left arm of the phase boundary (the binodal) denotes the saturation concentration *c*_*A*_ in the dilute phase, and the right arm of the binodal corresponds to the concentration *c*_*B*_ of the scaffold in *B*, which is the dense phase. The polyphasic linkage formalism describes how ligand binding modulates *c*_*A*_ for fixed solution conditions. The value of *c*_*A*_ in the presence of the ligand, designated as 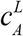, is determined by equalizing the chemical potential of the scaffold across the phase boundary. This yields the expression 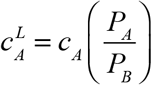 Here, *P*_*A*_ and *P*_*B*_ are the binding polynomials that quantify binding of the ligand to the scaffold in phases *A* and *B*, respectively. If *P*_*A*_ is greater than *P*_*B*_, then the ligand binds preferentially to the scaffold in phase *A*. Preferential binding of the ligand to the scaffold in the *A* phase will lead to an increase in 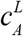 compared to *c*_*A*_, thus weakening the driving forces for phase separation of the scaffold (**Fig. S1A**, *SI Appendix*). Conversely, if *P*_*B*_ is greater than *P*_*A*_, then the ligand binds preferentially to the scaffold in the dense phase *B*. Accordingly, 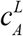 decreases when compared to *c*_*A*_. In this scenario, preferential binding of the ligand to the scaffold in the dense phase will enhance the driving forces for phase separation as evidenced by lowering of the threshold concentration to be crossed in order for the system to undergo phase separation (**Fig. S1B**, *SI Appendix*). In the absence of preferential binding, the ligand binds equivalently to the scaffold in both phases, implying that 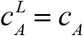 and the ligand behaves like a passive client that does not alter the phase equilibrium of the scaffold.

A weakness of the polyphasic linkage formalism is that it does not yield information regarding the specific features of ligands that lead to preferential binding to one phase over the other. Here, we remedy this using the *stickers*-and-*spacers* model (14, 21) to uncover rules for how precise control over scaffold phase behavior can be achieved through the preferential interactions of ligands. To uncover these rules, we use a coarse-grained linear polymer model that mimics well-known examples of scaffolds (1, 4, 8–10, 13, 22–24). We assess the effects of five types of ligands. These include two that were previously examined (25–27) and three new ligand types namely, monovalent ligands that interact with scaffold stickers, monovalent ligands that interact with scaffold spacers, and bipartite ligands that interact with scaffold stickers and spacers. Through the use of these models, we generate insights regarding the features of ligands that stabilize or destabilize condensate formation by scaffolds. Importantly, we: (a) determine how specific features of ligands contribute to modulating scaffold phase behavior, (b) provide mechanistic explanations for why ligands promote or destabilize scaffold phase behavior; and (c), quantify how the structure and concentration of the dense phase change upon ligand binding. Our work builds on key contributions reported recently that use patchy spherical particles as models for scaffolds and ligands (25–27). These studies have helped elucidate certain aspects of how low valence patchy ligands exert control over the phase behavior of high valence patchy scaffolds (27).

In addition to generating insights regarding ligand-mediated regulation of condensates, we highlight the importance of directly measuring how ligands affect phase boundaries of scaffolds. Such measurements, performed in live cells, are likely to pave the way for understanding how cells control condensate formation and dissolution via preferential binding of specific types of ligands at the right place and at the right time. Our analysis also shows that uncovering the modulatory effects of ligands cannot be achieved by measuring partition coefficients (*PCs*) of ligands. This is because the *PC* of a ligand is a convolution of many factors. Accordingly, high values of *PCs* for ligands do not have to mean a preferential binding to scaffolds in the dense phase, nor do they tell us anything about the modulatory effects of ligands on scaffold phase behavior.

## Results

### Coarse-grained model for examining the effects of different ligands on scaffold phase behavior

We deployed coarse-grained simulations using the lattice simulation engine LASSI (21) to understand how different types of ligands modulate the phase behavior of model linear multivalent macromolecules (details in *SI Appendix*). Linear multivalent macromolecules are represented by *stickers*, which are sites that drive phase separation, and *spacers*, which are sites interspersed between stickers that influence the interplay between phase separation and percolation (21, 22). Here, spacers can be implicit, in that they do not take up volume, or they can be explicit, in that they occupy volume on the lattice.

Previous studies have examined how the valence, sticker interaction strengths, and spacer excluded volumes of linear multivalent macromolecules effects scaffold phase behavior (4, 15, 22, 23). Accordingly, we focus here on a single type of scaffold molecule. In our model system, the scaffold molecule contains five sticker sites and two explicit spacer sites (**Fig. 2A**). The inclusion of explicit spacer sites allows for modeling the effects of ligands that interact directly with spacer sites. The specific instantiation of the *stickers*-and-*spacers* model used here helps ensure that: (1) the valence of spacer sites is less than the valence of sticker sites such that phase separation is driven mainly by sticker-sticker interactions regardless of ligand type; and (2) we observe robust phase separation in the concentration regime examined – an important design criterion given that increasing the number of explicit spacer sites can destabilize phase separation (22).

**Figure 2:**
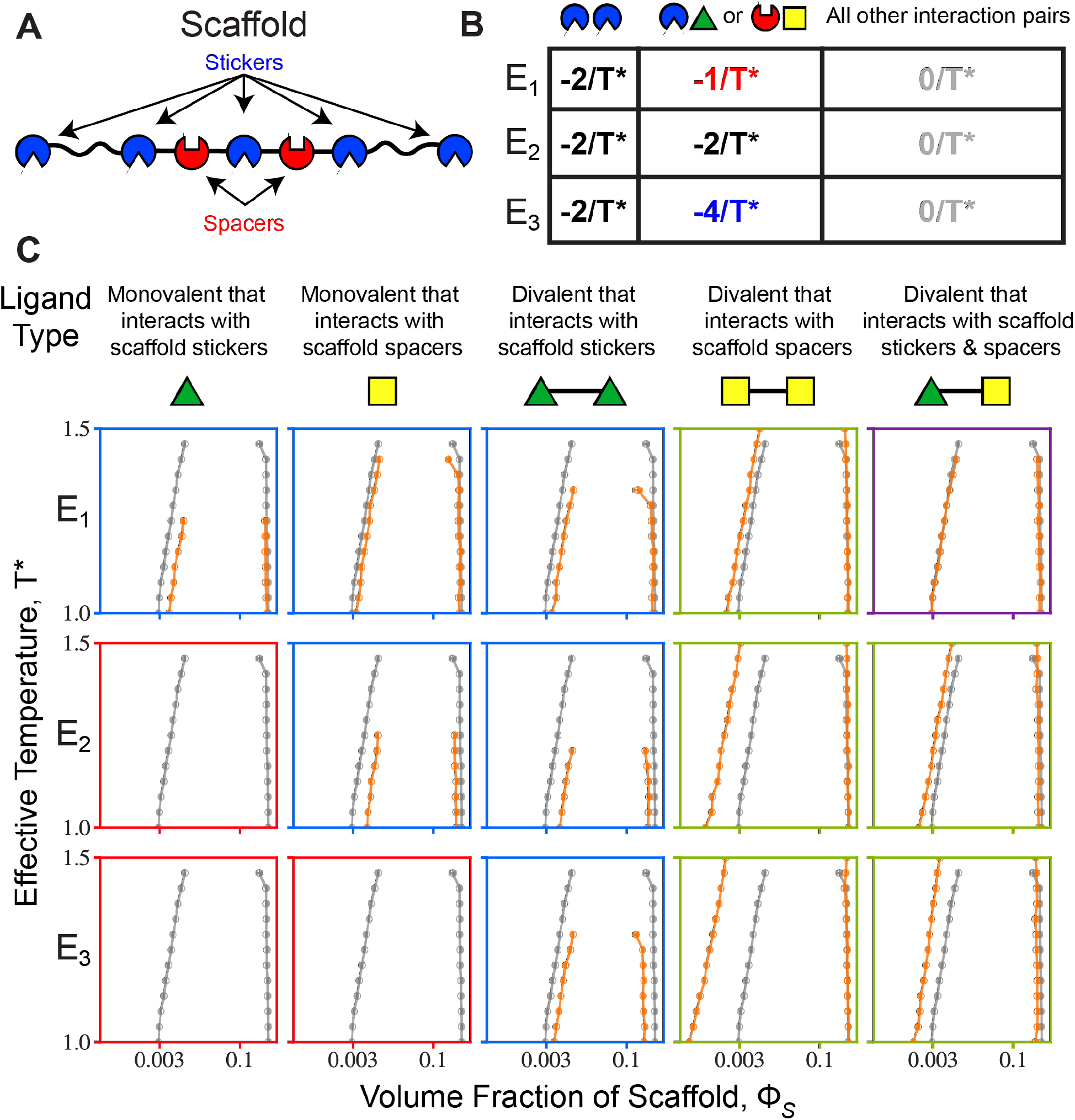
Effect of ligand types on scaffold phase behavior. (**A**) Schematic of the scaffold molecule used in coarse-grained simulations. The molecule has five sticker sites and two explicit spacer sites (the remaining two spacer sites are implicit). (**B**) Three energy scales were examined to assess how the different ligands modulate scaffold phase separation. (**C**) Binodals of the scaffold in the absence of ligand (grey) and in the presence of the given ligand (orange) for the three different energy scales, E_1_, E_2_, and E_3_. The bounding boxes for each case are color coded to summarize the effect of each ligand on scaffold phase separation: red – ligand binding abolishes phase separation, blue – ligand binding destabilizes phase separation, purple – ligand binding does not change phase separation, and green – ligand binding promotes phase separation.

To determine the effect of ligand type on scaffold phase behavior, we consider five different ligand types (**Fig. 2C**): (i) a monovalent ligand that interacts exclusively with scaffold stickers; (ii) a monovalent ligand that interacts exclusively with scaffold spacers; (iii) a divalent ligand that interacts exclusively with scaffold stickers; (iv) a divalent ligand that interacts exclusively with scaffold spacers; and (v) a bipartite divalent ligand that interacts with scaffold stickers and spacers. The only attractive interactions in the system are between pairs of sticker sites on scaffolds and between scaffold sticker or spacer sites and ligands. The details of the latter will depend on the type of ligand being considered. Additionally, each site can engage in only one interaction at a time. We consider three energy scales for the ligand-scaffold interaction (**Fig. 2B**): E_1_, the ligand-scaffold interaction is half that of the scaffold-scaffold interaction, E_2_, the ligand-scaffold interaction is equal to the scaffold-scaffold interaction, and E_3_, the ligand-scaffold interaction is double that of the scaffold-scaffold interaction. For each case, we performed five independent simulations as detailed in the *SI Appendix*.

### Quantifying the effects of different types of ligands on phase diagrams

The simulations allow us to probe the effects of ligand binding on the low and high concentration arms of scaffolds. **Fig. 2C** shows binodals for the scaffold without ligand and the impact of ligand binding on the binodals. In each case, we show results for a ratio of 0.23 for the ligand to scaffold sites. For this ratio, we observe four categories of ligand-modulated phase behaviors of scaffolds: (1) Ligand binding abolishes scaffold phase separation (phase diagrams with red bounding boxes in **Fig. 2C**), (2) ligand binding destabilizes scaffold phase separation (phase diagrams with blue bounding boxes in **Fig. 2C**), (3) ligand binding does not impact phase separation (phase diagrams with purple bounding boxes in **Fig. 2C**), and (4) ligand binding promotes phase separation (phase diagrams with green bounding boxes in **Fig. 2C**). Results for three additional ligand-to-scaffold ratios are shown in the *SI Appendix* (**Figs. S2-S4**), and the results are qualitatively similar to those shown in **Fig. 2C**.

Monovalent ligands destabilize or abolish scaffold phase separation, and this is true irrespective of whether they interact with sticker or spacer sites (columns 1 and 2 of **Fig. 2C**). Divalent ligands that interact directly with sticker sites on scaffolds also destabilize phase separation by competing with the sticker-sticker interactions that drive phase separation (column 3 of **Fig. 2C**). In contrast, phase separation is stabilized by divalent ligands that interact with scaffold spacer sites (column 4 of **Fig. 2C**). Bipartite divalent ligands that bind both sticker and spacer sites of scaffolds show an intermediate effect compared to the other two divalent ligands. They can promote phase separation at higher interaction strengths; however, this effect is weaker when compared to that of divalent ligands that bind only to spacer sites (column 5 of **Fig. 2C**).

### Quantifying the effects of different types of ligands on dilute phase concentrations of scaffolds

Experimental characterizations of full binodals are challenging, and accordingly, these measurements have been performed only for a small number of scaffold molecules (28–31). In contrast, it is easier to measure changes in saturation concentrations in the absence (4) and the presence of ligands (13, 32). To set up expectations regarding how saturation concentrations change, we analyze how 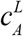 changes as a function of ligand concentration for each of the five ligand types (**Fig. 3**). At a given ligand concentration, the greater the difference between 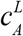 and *c*_*A*_, the greater the asymmetry in the preferential binding of the ligand to the scaffold in either the dense or dilute phase.

**Figure 3:**
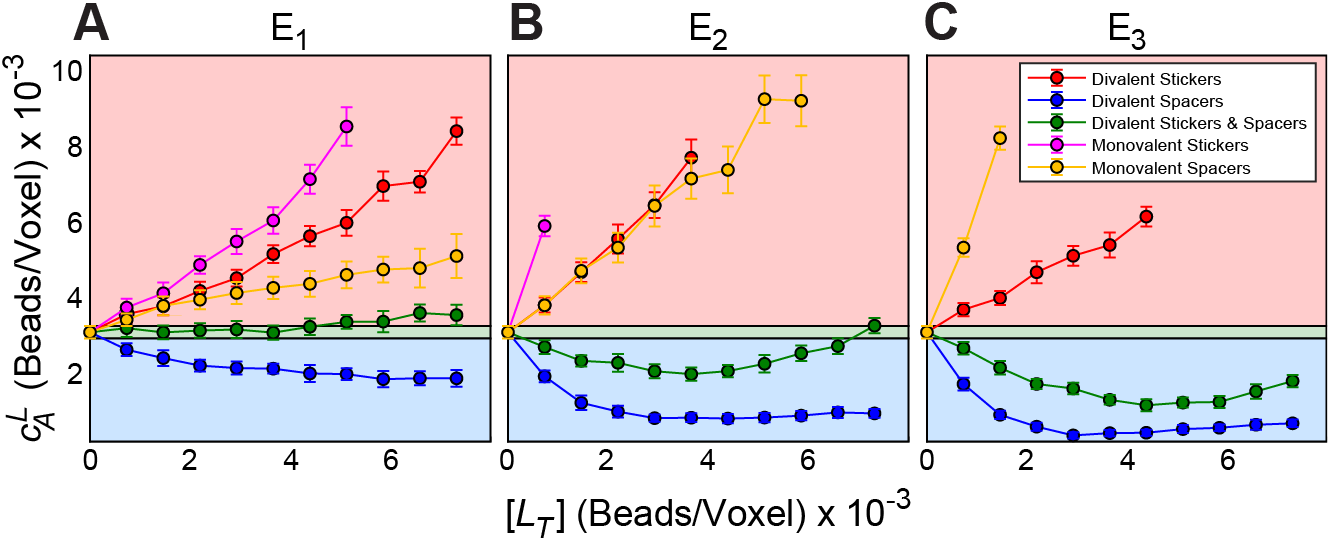
Changes to 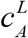 by each ligand type as a function of ligand concentration for different energy scales: E_1_, E_2_, and E_3_. Data are shown for *T*\* = 1.8. Shaded regions represent whether the ligand destabilizes (red), does not change (green) or promotes (blue) scaffold phase separation. Data are plotted only if the system undergoes phase separation, i.e., the width of the two-phase regime satisfies the criterion 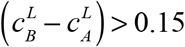

For all ligand concentrations and interaction strengths tested here, both monovalent ligands and the divalent ligand that interacts with scaffold stickers cause an increase in 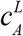 thereby destabilizing scaffold phase separation. The extent of destabilization increases monotonically as the energy scale is increased for the monovalent ligands (see magenta and orange traces in panels A-C of **Fig. 3**). However, the destabilizing effects of the divalent ligand that interacts with scaffold stickers changes non-monotonically with the energy scale. For a fixed ligand concentration, the extent of destabilization increases upon doubling the ligand-scaffold interaction energy from E_1_ to E_2_. However, the extent of destabilization then decreases upon further doubling of the ligand-scaffold interaction energy from E_2_ to E_3_. Multivalent ligands that bind to scaffold stickers tend to destabilize phase separation by competing with sticker-sticker interactions. However, at higher ligand-scaffold interaction strengths, the extent of destabilization can be reduced because the system now uses ligand-mediated crosslinks. Therefore, the interplay between ligand valence and the relative strengths of scaffold-scaffold versus ligand-scaffold interactions can be modulated to obtain non-monotonic changes to condensate stability.

The divalent ligand that interacts with scaffold spacers promotes scaffold phase separation as evidenced by the fact that 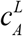 decreases vis-à-vis *c*_*A*_ for all ligand concentrations and interaction strengths tested here. We observe a weak non-monotonic trend in that 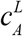 decreases and then increases with increasing ligand concentration. A similar effect has been reported in experiments that characterized the phase behavior of a poly-SH_3_-poly-PRM system in the presence of increased concentrations of the ligand heparin (26). For this system, the non-monotonic behavior was hypothesized to be due to electrostatic repulsions of heparin at high concentrations. Since charge effects are not included in our model, the simplest explanation is that the non-monotonic behavior results from a ligand concentration dependent interplay between scaffold-scaffold interactions being the only drivers of phase separation to some of these scaffold-scaffold interactions being competed out by ligand-scaffold interactions. The latter is a consequence of increased ligand concentration, which means that it becomes more likely to make ligand-scaffold interactions when compared to scaffold-scaffold interactions.

For the bipartite divalent ligand that interacts with sticker and spacer sites on scaffolds the ligand causes minimal changes to 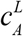 when compared to *c*_*A*_ for the lowest interaction strength, E_1_. However, there is a modest destabilization of scaffold phase separation at the highest ligand concentrations (**Fig. 3A**). At higher interaction strengths and low ligand concentrations, the ligand promotes phase separation, but to a weaker extent than the divalent ligand that interacts purely with spacer sites on scaffolds. Again, when the bipartite divalent ligand promotes scaffold phase behavior, we observe non-monotonic behavior in the dependence of 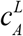 on ligand concentration. This non-monotonic behavior is a general feature of ligands that promote phase separation by preferentially binding to scaffolds in the dense phase.

The effects of bipartite ligands can be further tuned by changing the relative strength of the interaction between the site that interacts with scaffold stickers and the site that interacts with scaffold spacers (**Fig. S5**, *SI Appendix*). When the interaction strength with the scaffold spacer site is stronger than the interaction strength with the scaffold sticker site scaffold phase separation is promoted. In contrast, when the interaction strength with the scaffold sticker site is stronger than the interaction strength with the scaffold spacer site scaffold phase separation is destabilized. These results show that modulating the relative sticker versus spacer interaction strengths within a ligand provides an additional handle for modulating ligand mediated control of scaffold phase behavior.

### Impact of ligands on dense phase concentrations

We assessed how the scaffold and total dense phase concentrations in the presence of each of the ligand types change compared to the behavior of the scaffold alone (**Fig. 4**). The key observations are as follows: The scaffold concentration in the dense phase does not generally increase above the ligand-free case regardless of how ligand binding influences *c*_*A*_ (**Fig. 4A-C**). Ligands, irrespective of whether they enhance or weaken phase separation, will have a diluting effect on scaffolds within the dense phase and this effect increases with increasing ligand concentration (**Fig. 4A-C**). Dilution of the dense phase upon increasing ligand concentration has been observed experimentally for hnRNPA1 with BSA as a ligand (33). The extent of dilution depends on the interaction mode, interaction strengths, and ligand concentration. Specifically, binding to sticker sites on scaffolds has a greater effect on reducing the scaffold concentration in the dense phase, and this effect is increased as the interaction strength between the scaffold and ligand is increased. Further, ligands that do not change *c*_*A*_ can still modulate dense phase properties by reducing the scaffold concentration in the dense phase. The structural consequences, whereby ligands dilute the concentrations of scaffolds within the dense phase, follow from the requirement that the scaffolds have to accommodate ligands within condensates.

**Figure 4:**
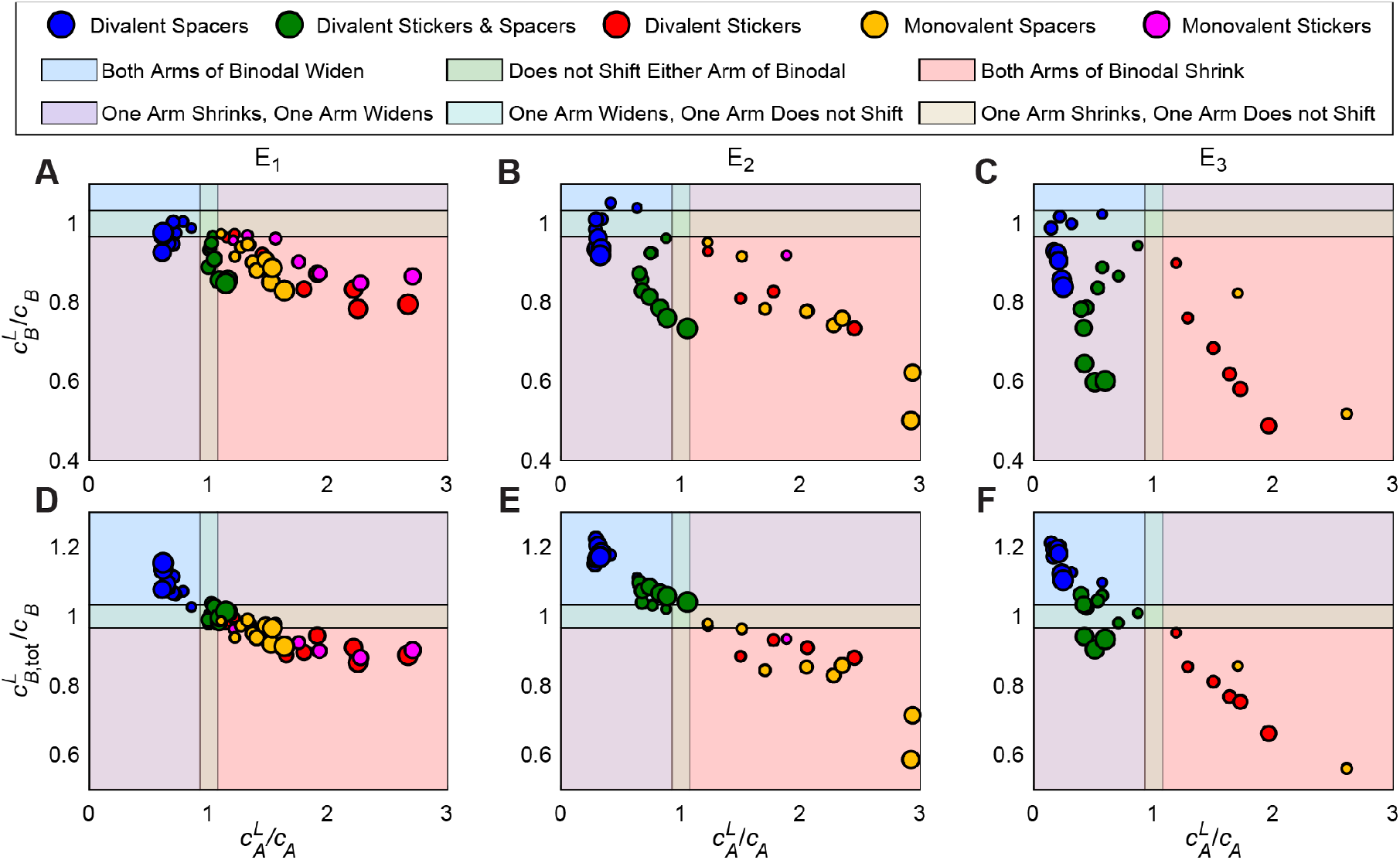
Change in dense phase versus dilute phase concentrations as a function of ligand type, interaction strength, and ligand concentration at T*≈1.08. The size of each point increases with ligand concentration. Data are plotted only if the system undergoes phase separation, i.e., the width of the two-phase regime satisfies the criterion 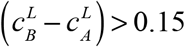

Next, we assessed the extent to which dilution of the scaffold concentration within condensates is compensated by an increase in ligand concentration. We quantified the total concentration of scaffolds *and* ligands in the dense phase for each of the different ligand types. The results, shown in **Figs. 4D-F**, may be summarized as follows: ligands that promote phase separation tend to increase the total dense phase concentration (blue); ligands that do not alter the driving forces for phase separation tend to maintain the total dense phase concentration (green); and ligands that destabilize phase separation tend to decrease the total dense phase concentration (red).

### Structural effects of ligands that bind preferentially to scaffolds in dilute versus dense phases

To uncover a “molecular” level understanding of the observations summarized above, we quantified site-to-site radial molecularity profiles, which we denote as *N*(*r*_*i*_) and define as the number of sticker or ligand occupied lattice sites in a shell that lies in the interval (*r*_*i*_, *r*_*i*_ +Δ) from each scaffold sticker site in the simulation volume. Details of how *N*(*r*_*i*_) profiles are calculated using pair distribution functions are provided in the *SI Appendix.*

Row A in **Fig. 5** shows scaffold sticker-to-sticker radial molecularity profiles plotted for *r* ≤ *L*/2, where *L* is the length of the side of the cubic simulation box. The short-range peak is a signature of phase separation (21). Destabilization of condensates leads to a ligand concentration dependent decrease and eventual abrogation of the first peak. This is seen in the left three panels of row A of **Fig. 5**. The monovalent ligands that bind only to spacer sites of scaffolds cause a dilution of sticker-to-sticker contacts. This is realized by enhancing the effective excluded volume of spacers and weakening the cooperativity of the inter-sticker crosslinks needed for driving phase separation (compare **Fig. 5C**, panels 1 and 3). Row B of **Fig. 5** shows the sticker-to-ligand radial molecularity profiles. For destabilizing ligands, a short-range peak is either present only at low ligand concentrations or non-existent as seen in the three left most panels of row B of **Fig. 5**. This is the result of ligand-mediated destabilization of condensates and the formation of sticker-ligand interactions in the dispersed one-phase regime.

**Figure 5:**
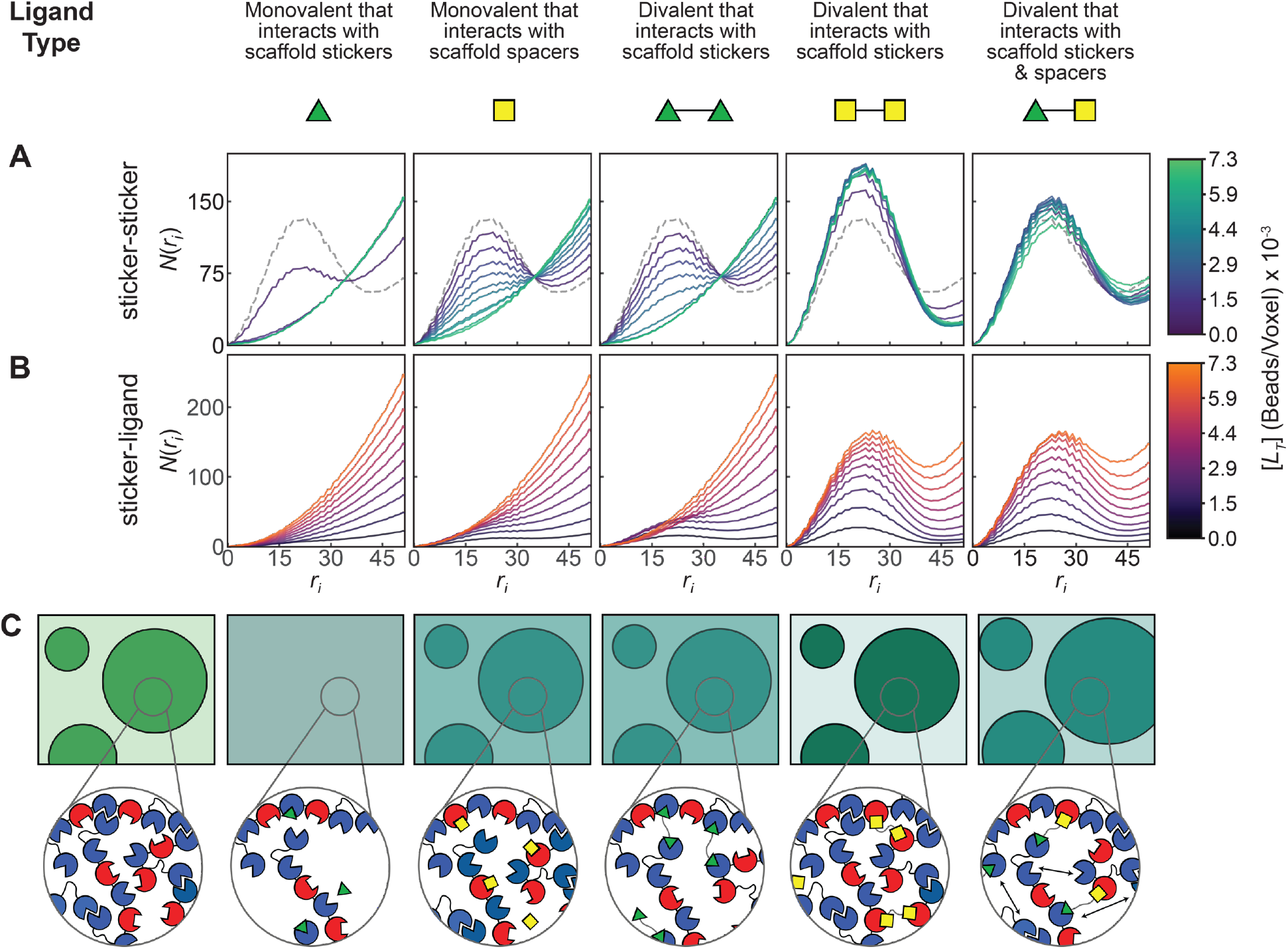
Analysis of site-to-site radial molecularity profiles. Data are shown for ligand-scaffold interaction strengths set to E_2_. **(A)** Sticker-to-sticker radial molecularity profiles quantifying how scaffold stickers organize around one another for different ligand concentrations and types. The dashed profiles are for ligand free cases. The presence of a peak in the profile is indicative of phase separation, and the heights of these peaks, which change with ligand concentration, quantify the effects of ligands on condensate stability. Monovalent ligands and divalent ligands that interact with stickers, weaken the sticker-sticker interactions, diluting stickers around one another, and thereby destabilize and / or dissolve condensates. **(B)** Sticker-to-ligand radial molecularity profiles that quantify the organization of ligands around scaffold sticker sites for different ligand types and ligand concentrations. The preferential accumulation of ligands around sticker sites in dense phases is evident in the growth of the height of the first peak with increasing ligand concentration. (**C**) Schematic summarizing the effects of ligands on the structural organization of scaffold stickers. The first box is in the absence of ligands. The insets for each box depict the various interactions occurring for that system. Blue sites denote scaffold stickers and red sites denote scaffold spacers. Scaffold sticker-sticker interactions are destabilized for the monovalent ligands and for the divalent ligand that binds scaffold stickers. The divalent ligand that binds scaffold spacers and the bipartite divalent ligand maintain scaffold sticker-sticker interactions and also provide additional networking interactions. These interactions for the bipartite ligand cause an increase in the correlation length between scaffold stickers within the dense phase.

For divalent ligands that bind directly to spacers and bipartite ligands that bind to spacers and stickers, we observe a maintenance of the first peak in both sets of radial molecularity profiles at all ligand concentrations (see right two panels in rows A and B of **Fig. 5**). The height of the first peak of the sticker-to-sticker radial molecularity profile shows non-monotonic behavior. The non-monotonic behavior can be explained by the increase in the height of the first peak in the sticker-ligand radial molecularity profile as ligand concentration increases (see right two panels in row B of **Fig. 5**). Accordingly, as ligand concentrations increase, ligand-mediated crosslinking become auxiliary drivers of scaffold phase separation.

The bipartite divalent ligands cause a dilution of scaffolds within the dense phase as shown in **Fig. 4B**. This can be explained by the radial molecularity profiles. There is a slight increase in the first peak of the sticker-to-sticker radial molecularity profile vis-à-vis the ligand-free case (last panel in **Fig. 5A**). This increase is followed by a decrease, rightward shift and widening of the first peak of the sticker-sticker profiles. Concurrently, the heights of the first peaks of ligand-to-sticker radial molecularity profiles increase monotonically and also exhibit rightward shifts. The inference is that scaffold sticker-sticker interactions are replaced by scaffold-ligand interactions thereby increasing the correlation length between scaffold stickers (last panel in **Fig. 5C**). This implies that at the highest ligand concentrations even though 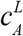 does not change relative to *c*_*A*_, the structural organization of scaffold stickers in the dense phase still changes due to interactions with the ligand. These alterations to the structural organization of scaffolds within condensates points to an additional regulatory function that ligands can exert over condensates. The impacts of ligands on structural organization of scaffold sites within condensates should be testable via suitable scattering experiments *in vitro* (34) or super resolution (35) based measurements in cells.

### Can partition coefficients provide information regarding preferential binding effects of ligands?

To answer this question, we calculated the *PCs* for the divalent ligands at *T*\*≈1.08. The ligand *PC* is defined as the concentration of the ligand in the dense phase divided by the concentration of the ligand in the dilute phase. **Figs. 6A** and **6B** show that there is a general trend for *PC*s to increase as ligands increasingly promote phase separation. However, when we examine a particular ligand concentration, we find that the *PC* for ligands that destabilize phase separation can be greater than *PC*s that correspond to ligands that promote phase separation (**Fig. 6C**). This behavior depends, at least partially, on the strength of the interaction between the ligand and the scaffold. These results suggest that the rank ordering of the *PC*s is not useful for discerning the effects of ligand on scaffold phase behavior.

**Figure 6:**
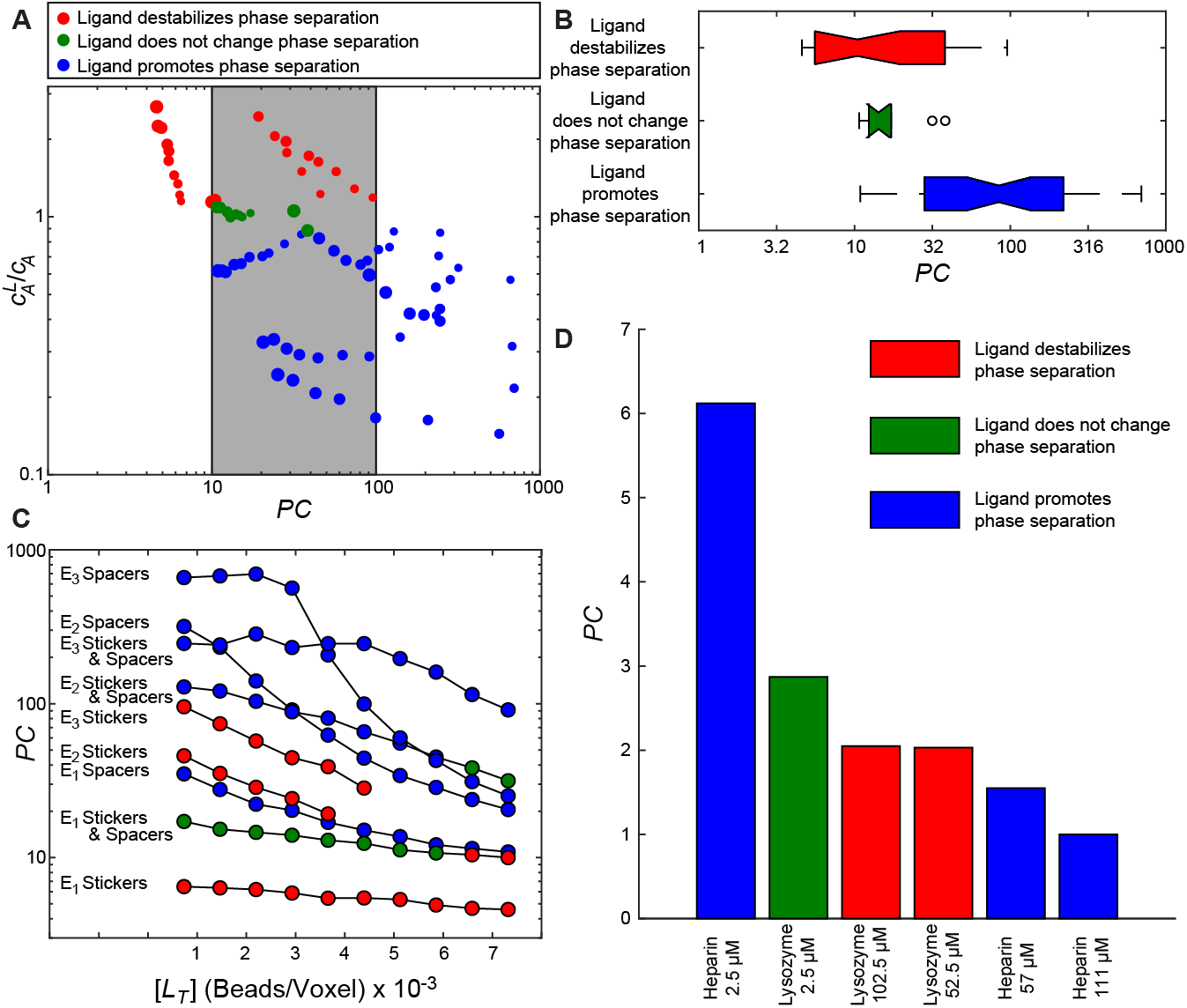
Comparing ligand partition coefficients to the modulation of the scaffold saturation concentration. (A) The changes in saturation concentration for all divalent ligands at *T*\*≈1.08 are plotted against the corresponding *PCs*. The size of each circle is proportional to the ligand concentration. The grey box denotes a region of *PC*s that is consistent with ligands that destabilize, do not change, or promote scaffold phase separation. Data are plotted only if the system undergoes phase separation, i.e., the width of the two-phase regime satisfies the criterion 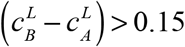 (B) Boxplots of the data presented in (A). (C) *PC*s as a function of ligand concentration for each of the divalent ligands and energy scales examined in this paper. (D) Parition coefficients measured by Ghosh et al. colored by the effect of the ligand on scaffold phase behavior at the same concentration. In all plots, blue implies the ligand promotes scaffold phase separation, green implies the ligand does not change the drive to phase separate, and red implies the ligand destabilizes scaffold phase separation.

Figs. 6A-B also show that there is a large range of ligand *PC* values that can correspond to all three modulatory effects of ligands on phase behavior. Specifically, we find that *PC* values spanning from 10 to 100 can correspond to ligands that destabilize, do not change, or promote scaffold phase separation. Additionally, all *PC*s are found to be greater than one. We also find that for a given ligand, *PC*s decrease with increasing ligand concentration regardless of the ligand modulatory effect on scaffold phase behavior (**Fig. 6C**). Together these results suggest that *PC*s are not a direct measure of preferential ligand binding. This is because *PCs* of ligands, unlike those of scaffolds, are a convolution of factors including ligand concentrations and interaction strengths of ligands for scaffold sites.

Ghosh et al., (26) recently reported a complete assessment of ligand *PC*s and their effects on scaffold phase behavior. Focusing on the poly-SH_3_:poly-PRM system, Ghosh et al., measured how lysozyme and heparin modulated the saturation concentration of poly-SH_3_:poly-PRM condensates. At high concentrations, lysozyme destabilizes the formation of poly-SH_3_:poly-PRM condensates. In contrast, heparin promoted the formation of poly-SH_3_:poly-PRM condensates at low and intermediate heparin concentrations, but destabilized condensate formation at high heparin concentrations. Additionally, Ghosh et al., measured *PC*s at three different ligand concentrations where they quantified the effects of ligands on condensate stability. **Fig. 6D** summarizes the key takeaways from these experiments. The bar color indicates the effect on condensate formation, where blue indicates promotion of phase separation, green indicates no change in phase behavior, and red indicates destabilization of phase separation. Both lysozyme and heparin show a decrease in *PC* as the ligand concentration is increased. Our simulation results are in agreement with these observations from experiments. Further, the data of Ghosh et al., show that *PC*s of ligands that promote phase separation can be lower than those of ligands that destabilize phase separation, even though the experiments were performed at similar bulk concentrations of ligands.

### Linkage theory (20) establishes that partition coefficients combine the contributions of preferential binding and local concentration effects

The impact of ligand binding on saturation concentrations is written in terms of binding polynomials *P*_*A*_ and *P*_*B*_ that quantify the binding of the ligand in question to the scaffold in phases *A* and *B*, respectively. A binding polynomial is the partition function of the ligand plus scaffold system and is a sum over the activities of all states in the system involving the scaffold relative to the free scaffold (36). We assume two types of systems, one where ligand binding to the scaffold is described by a first order polynomial in both phases and one where ligand binding to the scaffold is described by a second order polynomial in both phases. When binding can be described by a first order polynomial, 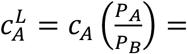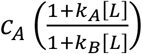, where [*L*] is the free ligand concentration and *k*_*A*_ and *k*_*B*_ are the association constants of the ligand to the scaffold in phase *A* and *B*, respectively. Likewise, when binding can be described by a second order polynomial, 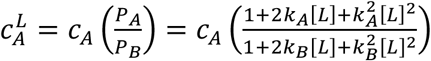.

In its simplest form, linkage theory assumes that the dense phase scaffold concentration does not change in the presence of ligand. From our coarse-grain simulations, we find that this assumption is only reasonable for systems with ligands that promote phase separation with a ligand to scaffold molecule ratio of less than two (**Fig. 4**). Therefore, we focused our analysis on systems with *k*_*B*_/*k*_*A*_ = 2, 4, 6, 8, and 10 and total ligand to scaffold concentration ratios spanning 0.25 to 1.75. The first criterion imposes preferential dense phase binding and hence the promotion of phase separation.

We examined the relationship between 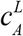 and *PC* by solving for the system of equations that describe each binding reaction and using the fact that the total scaffold concentration in phase *A* is given by 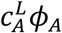, where *ϕ*_*A*_ is the volume fraction of phase *A* (see *SI Appendix* for details). In both phases the ligand can be free or bound, and the partitioning of free ligand between the two phases is governed by the relative volumes of each phase. Therefore, when binding is described by first order polynomials in both phases, it follows that 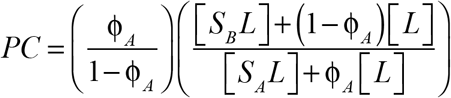, where [*S*_*A*_*L*] and [*S*_*B*_*L*] are the concentrations of the bound scaffold in phase *A* and *B*, respectively. Likewise, when binding is described by second order polynomials in both phases, 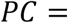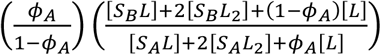, where [*S*_*A*_*L*_2_] and [*S*_*B*_*L*_2_] are the concentrations of the scaffold bound by two ligands in phase *A* and *B*, respectively.

**Fig. 7** shows how 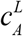 and *PC* change as a function of total ligand to scaffold ratio, *k*_*B*_/*k*_*A*_, and whether binding is first or second order. We set *c*_*A*_ = 1 μM, *c*_*B*_ = 19 μM, and the total scaffold concentration, [*S*_*T*_] = 10 μM. As designed, all systems show a decrease in 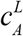 compared to *c*_*A*_, implying phase separation is promoted upon ligand binding (**Fig. 7A**). Consistent with our coarse-grained simulation results and the results of Ghosh et al., we observe that *PCs* decrease and approach one at high ligand concentrations for all systems (**Fig. 7B**). However, the slopes and details of how 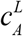 and *PC* change with ligand concentration vary depending on the binding mode.

**Figure 7:**
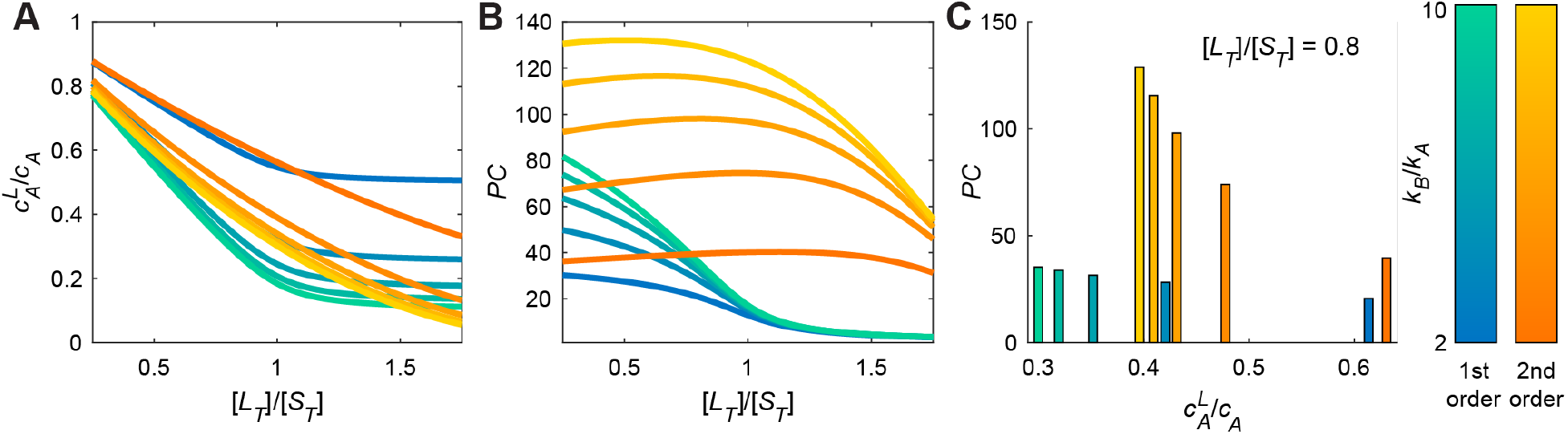
Theory shows how saturation concentrations and partition coefficients change as a function of ligand concentration for ligands with different binding association constants and binding modes. Ligands that bind as described by a first order polynomial are shown in the blue to green color scale and ligands that bind as described by a second order polynomial are shown in the orange to yellow color scale. (A) The change in scaffold saturation concentration as a function of the ratio of the total ligand concentration to the total scaffold concentration. (B) The partition coefficient as a function of the ratio of the total ligand concentration to the total scaffold concentration. (C) *PC* as a function of the change in the scaffold saturation concentration for a ligand to scaffold ratio of 0.8.

**Fig. 7C** examines the relationship between 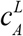 and *PC* at a ligand to scaffold ratio of 0.8. Although *PCs* decrease with increasing 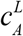 for a given binding mode, we find that there is a non-monotonic relationship between 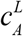 and *PC* when both binding modes are considered. In most cases, we will not have *a priori* information regarding the binding mode a ligand. This highlights the problem that even for ligands that promote phase separation, rank ordering *PC*s of ligands does not provide a sorting of ligands by the degree of their impact on promoting phase separation.

## Discussion

The *stickers*-and-*spacers* formalism (27, 37–40) allows us to uncover key features of ligands that destabilize or stabilize phase separation via preferential binding to scaffolds in the dilute versus dense phase, respectively. Overall, our findings are as follows: Monovalent ligands weaken phase separation either by reducing the overall valence of the scaffold when they interact directly with sticker sites or by enhancing the effective excluded volume of spacers and weakening the cooperativity of the inter-sticker crosslinks that is needed for driving phase separation when they interact directly with spacer sites. Divalent ligands that bind to scaffold sites weaken phase separation by competing directly with inter-sticker interactions. In contrast, divalent ligands that bind to spacer sites enable additional networking of multivalent scaffold molecules by serving as crosslinkers, thereby promoting phase separation. This shows that ligands can lower the saturation concentrations for scaffolds, a finding that is important in light of an ongoing debate about the relevance of phase separation *in vivo*, especially at endogeneous expression levels (18).

Our findings, and those of others (27), imply that scaffolds can undergo ligand-mediated phase separation even if the endogeneous concentration of the scaffold is below its intrinsic saturation concentration. This feature is likely to be amplified by the collective contributions of networks of ligands, providing they are multivalent (8–10). Finally, bipartite divalent ligands that bind both stickers and spacers within the scaffold can modulate scaffold phase behavior in either direction depending on the relative interaction strengths of the ligand with the stickers and spacers of the scaffold. Ligand-modulation of condensate stability may be thought of as being another component of *heterotypic buffering*, a concept recently introduced to describe how the interplay between homotypic and heterotypic scaffold-scaffold interactions regulates scaffold phase behavior *in vivo* (16).

We find that the concentrations of scaffolds within the dense phase stays similar to that of the unliganded case or decreases in the presence of ligand. This is true irrespective of whether or not the ligand binds preferentially to the scaffold in its dense or dilute phase. For preferential binding to the scaffold in the dense phase, 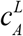 decreases, and the scaffold concentration in the dense phase 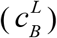 generally decreases when compared to *c*_*B*_. This helps accommodate ligands that bind preferentially to scaffold sites in the dense phase. Conversely, for preferential binding to the scaffold in the dilute phase, 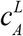 increases and 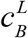 decreases compared to *c*_*B*_. This dilution of scaffold within the dense phase derives from weakening the driving forces for phase separation.

We also focused on the important question of how one might quantitatively assess the contributions of ligands as modulators of condensate formation and dissolution *in vivo* or *in vitro*. One approach would be to measure *PCs* of ligands since they quantify the enrichment or depletion of ligands in condensates (41). However, we show that *PCs* of ligands are convoluted quantities that do not provide direct assessments of the effects of ligands as modulators of scaffold phase behavior. Instead, the simplest approach would be direct measurements of scaffold concentrations in the dilute and dense phases as a function of ligand concentration (32).

It is worth noting that we have made the simplifying assumption that a scaffold will be defined by a fixed saturation concentration. However, this is only true if homotypic interactions among scaffold molecules are the primary drivers of phase separation (6, 21). If condensates form via a combination of homotypic and heterotypic interactions (6, 21, 42), then the network of these interactions (7, 12) and hence a combination of scaffold concentrations will determine the location of the phase boundary. In this scenario, one would have to measure the effects of ligands on the location of the phase boundary, governed jointly by the concentrations of all scaffold molecules that drive phase separation. This requires measuring the concentrations of more than one scaffold molecule while titrating the concentration of the ligand in question. Further complexities will arise as we consider how a set of distinct ligands impact the phase behavior of condensates that are governed by a network of homotypic and heterotypic interactions of scaffold molecules.

It is also known that the driving forces for phase separation can be modulated by anchoring scaffolds to surfaces (43) or via physical interactions with soft surfaces in cells (44). For example, Morin et al., (44) showed that preferential interactions of a pioneer transcription factor KLF4 with the surface of DNA can lower the threshold concentration for phase separation. Morin et al., used the Brunauer-Emmett-Teller theory (45) for multilayer adsorption to explain their results. Taken together, it follows that preferential binding of multivalent ligands to scaffolds in dense phases, this work and that of Ghosh et al., (26) and Espinosa et al., (27), and adsorption of scaffolds to surfaces (46, 47) as shown by Morin et al., (44) can lead to similar effects in terms of lowering the saturation concentrations of scaffold macromolecules. Therefore, we propose that the stabilization of condensate formation via surface interactions derives from preferential adsorption of dense phases through spacer-surface interactions and / or enhancing of sticker-sticker crosslinks. If this proposal is valid, then it follows that a unified theory is likely achievable for describing how the stabilities of condensates are impacted by the bulk phase concentrations of preferentially binding ligands and the surface features of soft interfaces.

Finally, our work suggests that the effects of small molecules (ideally, multivalent ligands) on 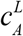 can be used as part of a chemical biology toolkit (48) to infer the features and internal organization of scaffolds within condensates. For example, if molecules with certain chemical structures destabilize condensates, then one can infer that a complementary interaction motif in the scaffold is primarily accessible in the dilute phase and thus may be involved in driving phase separation. These inferences have the potential to enable the design of small molecules that modulate scaffold phase behavior in prescribed ways.

## Materials and Methods

Full details of the LASSI simulations and corresponding analyses are given in the *SI Appendix*.

## ACKNOWLEDGMENTS

This work was supported by grants from the US National Institutes of Health (5R01NS056114, 1R01NS089932) and Dewpoint Therapeutics Inc. (through a sponsored research agreement with Washington University in St. Louis). We thank Megan Cohan, Mina Farag, Anthony Hyman, Alex Holehouse, Diana Mitrea, Mark Murcko, Michael Rosen, Ammon Posey, and J. Paul Taylor for helpful discussions.

