## Supplemental Material for "Ligand Effects on Phase Separation of Multivalent Macromolecules"

**This PDF file includes:**

Supplementary text
Figures S1 to S5
Tables S1 to S2
SI References

**Supplementary Information Text**

**Simulation Details.** To test how ligands affect the phase boundaries of macromolecules we used the lattice simulation engine LASSI (1). To the open-source version that we have made available on GitHub, we added analysis routines to enable the calculation of specific quantities that are unique to the current work. Note that the simulation engine itself is the same as the open-source one and all of the results presented in this work can be reproduced using the open-source version of LASSI that we have made available.

LASSI is designed for simulations of phase transitions of polymers on a lattice using Monte Carlo (MC) simulations. In the current simulations, 2000 chains of the macromolecular scaffold were used for every solution condition. To only have inter-molecular interactions, the scaffold was divided into two sets comprised of 1000 chains where each set had different sticker and spacer types. The total set of ligand numbers sampled corresponds to 400, 800, …, 4000 where for the monomer ligands the numbers were doubled to keep the ligand-site number and concentration consistent. Periodic boundary conditions were used, with a box size of $L$ = 103 lattice units. Each bead can only form one physical bond where bonds are stochastically formed in a way that preserves detailed balance over the course of the simulations. Mathematical details regarding the sampling techniques used are as prescribed in the original work on LASSI (1).

For any given solution condition (simulation temperature and combination of scaffold and ligand concentration), all molecules are randomly placed on the lattice and site-overlap is strictly forbidden. All interactions, other than site-overlap, are then turned off between the molecules. To accelerate the formation of one droplet for the dense phase, an external biasing potential is then applied to every bead which has the form $V(r) \sim (\vec{r} - \vec{r}_0)^2$ which serves as a means to push molecules towards the center of the simulation box. After $t_c = 5 \times 10^6$ MC sweeps, interactions between the molecules are turned on. The system is then simulated for $t_T = 2.5 \times 10^9$ MC sweeps for what we shall define as the first cycle.

Initially, the system is exponentially cooled down to $T_0 \equiv T^* = 1$, where the external potential now has the form $V(r,t) \sim e^{-4\frac{t}{t_c}}(\vec{r} - \vec{r}_0)^2$. When $T - T_0 \leq 0.005$, at $t_V \approx 7 \times 10^6$ we turn off the external potential ($V(r,t) = 0$). This results in the formation of only one large condensate provided the solution conditions correspond to the two-phase regime. The system is then run up till $t_T$. This concludes the first cycle.

The temperature of the system is then discontinuously increased by $\Delta T^* \approx 0.042$, and the system is simulated for $t_T$ MC sweeps, corresponding to the second cycle. 25 cycles are performed in total which sample $T^* = 1$ to $T^* = 2$.

5 independent sets of simulation are performed per solution condition using the following relative frequencies for the MC moves:

| | |
|---|---|
| Local | 3000 |
| Bond Rotation | 1500 |
| Co-local | 1500 |
| Multi-local | 1500 |
| Chain Translation | 500 |
| Chain Pivot | 500 |
| Double-pivot | 500 |
| Limited Cluster | 10 |
| Large Cluster | 1 |

Additionally, 5 independent runs are performed per solution condition. Averages over the requisite quantities defined below are performed over the last half of each cycle. Lastly, note that statistically similar results can be obtained without the initial external potential, but the simulations need to be about 10 times longer. For more details about the MC engine in LASSI, see (1).

**Constructing Binodals.** For every solution condition tested, we directly compute the co-existing densities via density-profiles referenced from the center-of-mass (COM) of the system. The density-profile is generated by computing a number histogram, $H(r_i)$, from the COM of the system with a bin-width of $0.25$, up to $r = \frac{\sqrt{3}L}{2}$, which corresponds to the maximal distance in a cubic box with periodic boundaries. To normalize the number histograms, we explicitly calculate the number histogram, $H_0(r_i)$, of lattice sites for a cubic box of dimensions $L = 103$ lattice-units with periodic boundaries, given a bin-width of $0.25$. Thus, the density profile is $\rho(r_i) = \frac{H(r_i)}{H_0(r_i)}$. To calculate $c_B$, we average over the first 7 bins excluding $r_0$, and for $c_A$ we average over the last 25 bins. For the binodals shown in **Fig. 2** and **Figs. S2-S4** only the co-existing densities are plotted where the densities are statistically different from each other.

**Generating Radial Molecularity Profiles.** For each simulation, we compute the radial molecularity profiles using radial pair-number histograms. The radial pair-number histograms are generated using the standard procedure where assuming isotropy, the radial singlet-number histogram, $H^{(1)}(r_i)$, is computed for a single bead. This radial histogram is then computed for every bead in the system where the sum over all beads then gives us the radial pair-number histogram $H^{(2)}(r_i)$ for the system. A bin-width of $1.0$ is used, for smoothness, and the histogram is computed up to $r = \frac{\sqrt{3}L}{2}$. Since we can track the bead-type of every bead, we can compute this pair-number histogram for every possible type-pair. To normalize the distributions, we divide the histogram by the total number of unique possible pairs for a given set of types being considered. For each type-pair, the number of pairs is $N_{ij} = N_i(N_j - \delta_{ij})/(1 + \delta_{ij})$ where $N_{i,j}$ represents the number of beads of type $i, j$ and $\delta_{ij}$ is Kronecker delta function. In the main-text, we focus on two sets: sticker-sticker and (scaffold)-sticker-ligand. Suppose that the sticker bead-types are $1$ and $2$, and that the ligand bead-types are $3$ and $4$. Then for the sticker-sticker set we have $1$-$2$ pairs only, while for the second set we have the sum of $1$-$3$, $1$-$4$, $2$-$3$ and $3$-$4$ pairs. To compute the radial molecularity profiles from these normalized histograms, we multiply each histogram by the total number of beads, $\widetilde{N}$, of the second index in the set. For the sticker-sticker set we multiply by the number of sticker beads of either type, $\widetilde{N} = 5000$. For the (scaffold)-sticker-ligand set we multiply by the total number of ligand beads at a given concentration $\widetilde{N}_m = 800\,m$ where $m$ is the concentration index. Therefore, the sum over each radial molecularity histogram is the number of sticker beads of either type for the first set and the number of ligand beads for the second set.

**Constructing Binding Polynomials.** We shall assume that the scaffold (*S*) has *n* independent ligand (*L*) binding sites in the dilute phase. The reaction equations for this process are given by:

$$[S] + i[L] \Leftrightarrow [SL_i]$$

where *i*=0,1,2,…*n*, $[SL_i]$ is the concentration of scaffold with *i* ligands bound, and $[S]$ and $[L]$ are the free scaffold and ligand concentrations, respectively. Let $k_j$ be the association constant for site *j*. Then, the concentration of the scaffold bound to a single ligand is:

$$[SL] = [S_1L] + [S_2L] + \cdots + [S_nL]$$

$$[SL] = k_1[S][L] + k_2[S][L] + \cdots + k_n[S][L]$$

where $[S_1L]$ denotes a single ligand bound to site 1 on the scaffold. Thus, the cumulative association constant for the scaffold bound to a single ligand is:

$$\beta_1 = \sum_{j=1}^{n} k_j = \frac{[SL]}{[S][L]}.$$

In general, the cumulative association constants are given by:

$$\beta_i = \frac{[SL_i]}{[S][L]^i}.$$

The binding polynomial is the summation of the concentration of all states in the system involving the scaffold relative to the free scaffold,

$$P = \sum_{i=0}^{n} \frac{[SL_i]}{[S]}.$$

Rewriting *P* in terms of cumulative associations constants, we get

$$P = \sum_{i=0}^{n} \beta_i [L]^i.$$

In the case where all binding sites are independent and identical, $k = k_1 = k_2 = k_n$,

$$\beta_i = \left(\frac{n!}{(n-i)!\,i!}\right) k^i$$

(2) and

$$P = (1 + k[L])^n.$$

**Connecting Linkage Theory to Partition Coefficients.** We shall consider an aqueous solution with a single type of scaffold that separates into two distinct phases. We denote the dilute phase as *A* and the coexisting dense phase as *B*. For a system where ligand binding to the scaffold is described by a first order polynomial in both phases, ligand binding is given by:

$$[S_A] + [L] \rightleftharpoons [S_A L]$$
$$[S_B] + [L] \rightleftharpoons [S_B L]$$

Here, $[L]$, $[S_A]$, $[S_B]$, $[S_AL]$ and $[S_BL]$ are respectively the concentrations of free ligand, free scaffold in phase *A*, free scaffold in phase *B*, bound scaffold in phase *A*, and bound scaffold in phase *B*. The binding polynomial in the dilute phase is given by:

$$P_A = \frac{[S_A]+[S_A L]}{[S_A]} = 1 + k_A[L];$$

Here, $k_A$ is the association constant in phase *A*. Likewise, the binding polynomial in the dense phase is $P_B = 1 + k_B[L]$, where $k_B$ is the association constant in the dense phase, *B*. Accordingly, it follows that: $c_A^L = c_A\left(\frac{P_A}{P_B}\right) = c_A\left(\frac{1+k_A[L]}{1+k_B[L]}\right)$. It should be noted that this equation may not be valid for high concentrations of free ligand because it ignores non-idealities due to interactions of ligands with themselves. Additionally, it assumes that $c_B$ does not change upon addition of the ligand.

To assess how $c_A^L$ and $PC$ are related we must solve the system specific set of equations that describe each binding reaction. Here, the system is defined by the following set of equations:

$$k_A = \frac{[S_A L]}{[S_A][L]},\ k_B = \frac{[S_B L]}{[S_B][L]},$$

$$[S_{A,T}] = [S_A] + [S_A L], \qquad [S_{B,T}] = [S_B] + [S_B L],$$

$$\text{and } [L_T] = [L] + [S_A L] + [S_B L].$$

Here, $[S_{A,T}]$ and $[S_{B,T}]$ are the total scaffold concentrations in phases $A$ and $B$, respectively, that are calculated using the total system volume $V$. Likewise, $[L_T]$ is the total ligand concentration in the system. The total scaffold concentration is given by:

$$[S_T] = \frac{n_A + n_B}{V_A + V_B} = \frac{n_A + n_B}{V} = [S_{A,T}] + [S_{B,T}].$$

Here, $n_A$, $n_B$ are the numbers of scaffold molecules within and $V_A$, $V_B$ are the volumes of phases $A$ and $B$, respectively; $V$ is the total volume of the system. Accordingly, the saturation concentration $c_A^L = \frac{n_A}{V_A}$ is related to $[S_{A,T}]$ as follows:

$$[S_{A,T}] = \left(\frac{n_A}{V}\right)\left(\frac{V_A}{V_A}\right) = c_A^L\left(\frac{V_A}{V}\right) = c_A^L \phi_A,$$

$$\text{and } [S_{B,T}] = [S_T] - [S_{A,T}] = [S_T] - c_A^L \phi_A.$$

Here, $\phi_A$ is the volume fraction of phase $A$. It follows that the system of equations can be rewritten in terms of $c_A^L$:

$$k_A = \frac{[S_A L]}{[S_A][L]},\ k_B = \frac{[S_B L]}{[S_B][L]},$$

$$[S_{A,T}] = [S_A] + [S_A L] = c_A^L \phi_A, \qquad [S_{B,T}] = [S_B] + [S_B L] = [S_T] - c_A^L \phi_A,$$

$$\text{and } [L_T] = [L] + [S_A L] + [S_B L].$$

By solving this system of equations, we can determine the ligand $PC$ given by:

$$PC = \left(\frac{\phi_A}{1-\phi_A}\right)\left(\frac{[S_B L] + (1-\phi_A)[L]}{[S_A L] + \phi_A [L]}\right).$$

Following the same procedure, for a system where ligand binding to the scaffold is described by a second order polynomial in both phases, the system of equations that describe each binding reaction is given by:

$$k_A = \frac{[S_A L]}{2[S_A][L]},\ k_B = \frac{[S_B L]}{2[S_B][L]},$$

$$k_A^2 = \frac{[S_A L_2]}{[S_A][L]^2},\ k_B^2 = \frac{[S_B L_2]}{[S_B][L]^2},$$

$$[S_{A,T}] = [S_A] + [S_A L] + [S_A L_2] = c_A^L \phi_A, \qquad [S_{B,T}] = [S_B] + [S_B L] + [S_B L_2] = [S_T] - c_A^L \phi_A,$$

$$\text{and } [L_T] = [L] + [S_A L] + 2[S_A L_2] + [S_B L] + 2[S_B L_2].$$

Here, $[S_A L_2]$ and $[S_B L_2]$ are the concentrations of scaffolds with two ligands bound in phase $A$ and $B$, respectively. Furthermore, when ligand binding can be described by a second order polynomial, $c_A^L = c_A\left(\frac{P_A}{P_B}\right) = c_A\left(\frac{1+2k_A[L]+k_A^2[L]^2}{1+2k_B[L]+k_B^2[L]^2}\right)$ and $PC = \left(\frac{\phi_A}{1-\phi_A}\right)\left(\frac{[S_B L]+2[S_B L_2]+(1-\phi_A)[L]}{[S_A L]+2[S_A L_2]+\phi_A[L]}\right)$.

**Fig. S1.** Illustration of polyphasic linkage. Scaffolds undergo phase separation and form two distinct phases: the scaffold-deficient phase (red), *A*, and the scaffold-rich phase (blue), *B*. The yellow region delineates the two-phase regime of the scaffold. The phase diagram is drawn in the two-parameter space of total scaffold concentration, $[S_T]$, and effective interaction strength among scaffold molecules. For a given value of the interaction strength along the ordinate, the left arm of the binodal represents the saturation concentration, $c_A$, and the right arm of the binodal denotes the concentration of the scaffold in the dense phase, $c_B$ (grey circles). Polyphasic linkage describes how $c_A$ changes by way of preferential binding of the ligand across the phase boundary and this is denoted as $c_A^L$. (A) $c_A^L > c_A$ if the ligand preferentially binds to the scaffold in the dilute phase; (B) $c_A^L < c_A$ if the ligand preferentially binds to the scaffold in the dense phase.

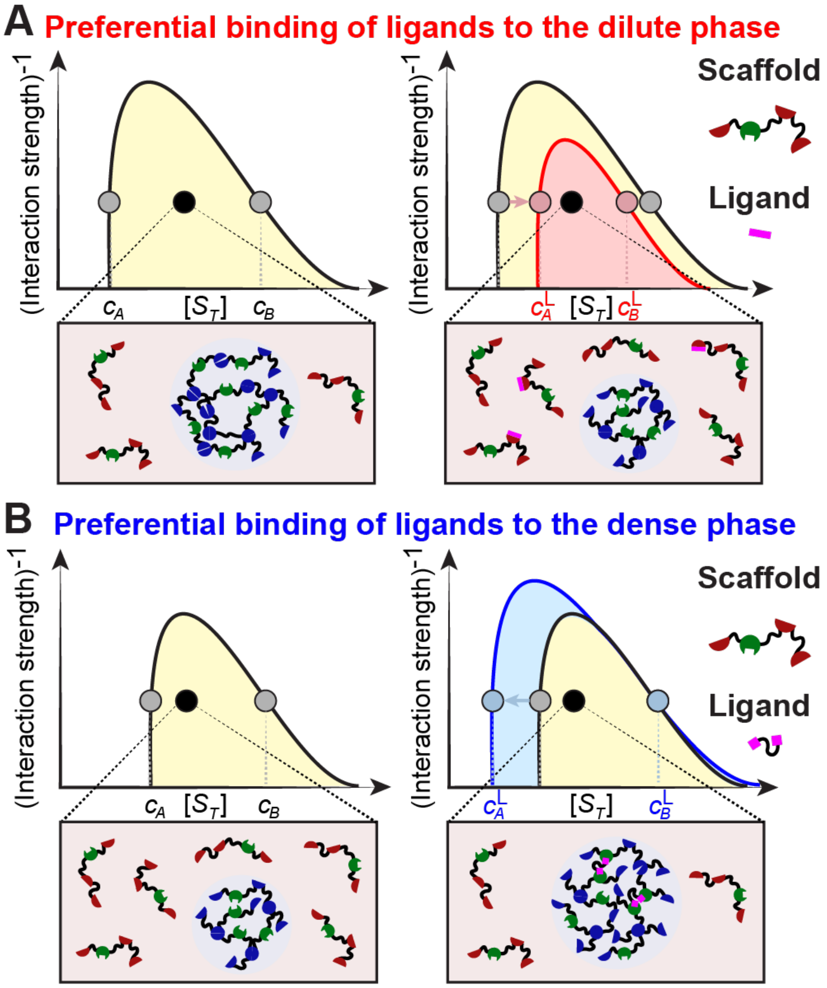

**Fig. S2.** Effect of ligand types on scaffold phase behavior for a ratio of 0.11 ligand to scaffold sites. Binodals of the scaffold in the absence of ligand (grey) and in the presence of the given ligand (orange) for the three different energy scales, $E_1$, $E_2$, and $E_3$. The bounding boxes for each case are color coded to summarize the effect of each ligand on scaffold phase separation: red – ligand binding abolishes phase separation, blue – ligand binding destabilizes phase separation, purple – ligand binding does not change phase separation, and green – ligand binding promotes phase separation.

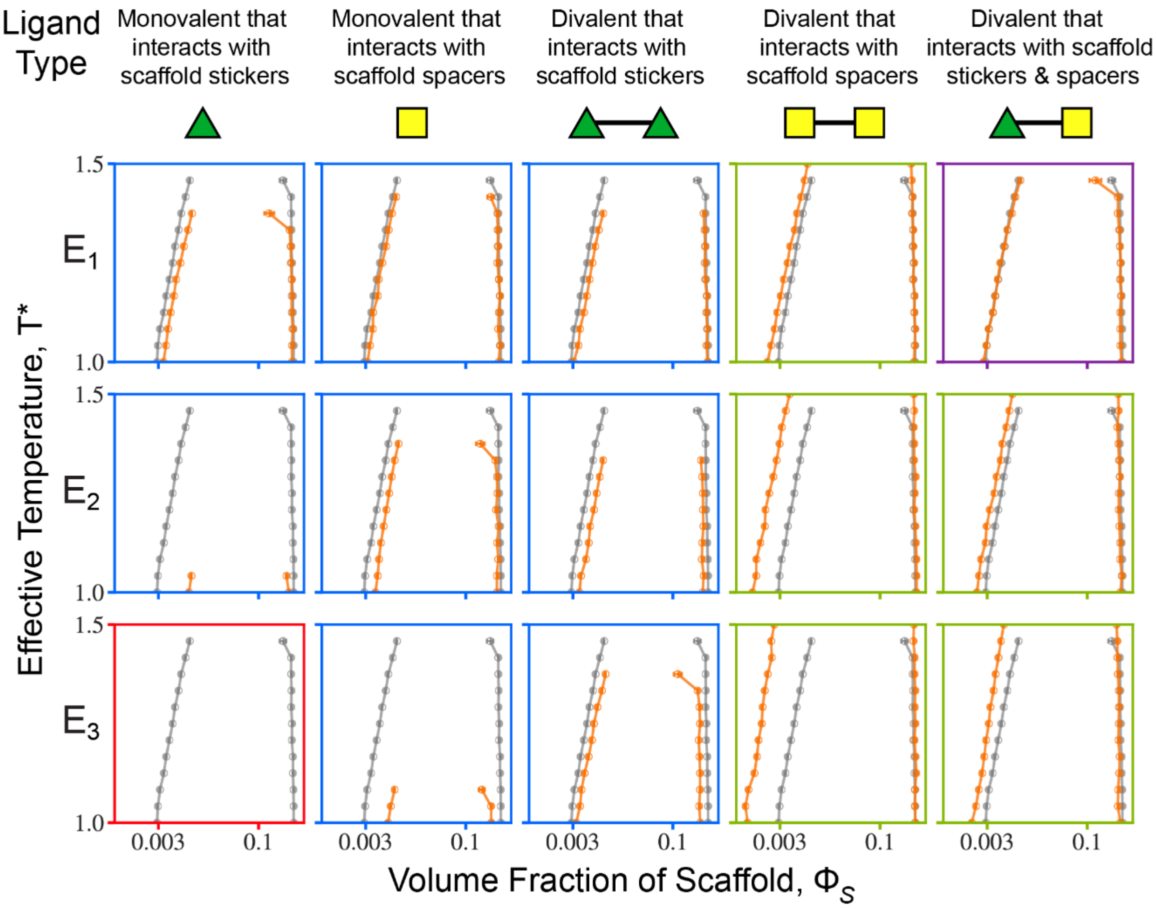

**Fig. S3.** Effect of ligand types on scaffold phase behavior for a ratio of 0.34 ligand to scaffold sites. Binodals of the scaffold in the absence of ligand (grey) and in the presence of the given ligand (orange) for the three different energy scales, $E_1$, $E_2$, and $E_3$. The bounding boxes for each case are color coded to summarize the effect of each ligand on scaffold phase separation: red – ligand binding abolishes phase separation, blue – ligand binding destabilizes phase separation, purple – ligand binding does not change phase separation, and green – ligand binding promotes phase separation.

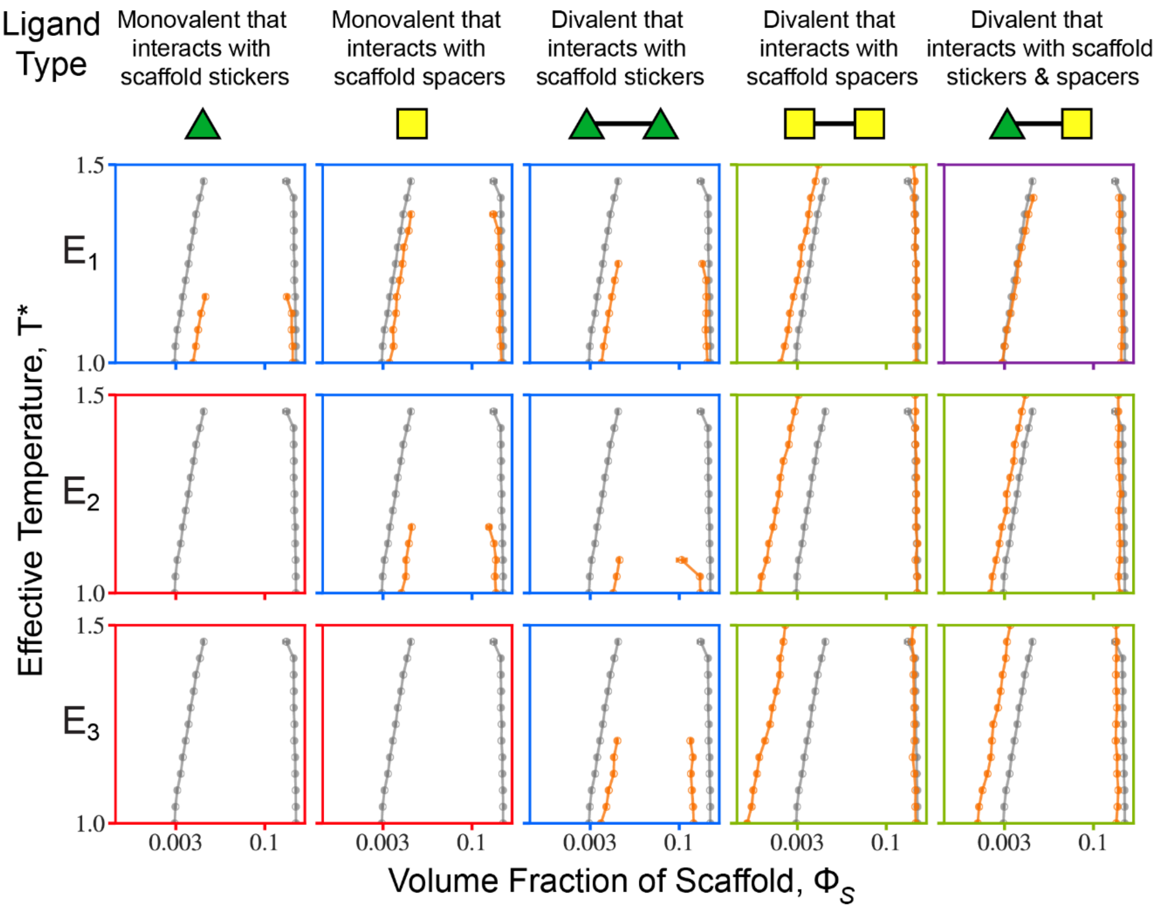

**Fig. S4.** Effect of ligand types on scaffold phase behavior for a ratio of 0.46 ligand to scaffold sites. Binodals of the scaffold in the absence of ligand (grey) and in the presence of the given ligand (orange) for the three different energy scales, $E_1$, $E_2$, and $E_3$. The bounding boxes for each case are color coded to summarize the effect of each ligand on scaffold phase separation: red – ligand binding abolishes phase separation, blue – ligand binding destabilizes phase separation, purple – ligand binding does not change phase separation, and green – ligand binding promotes phase separation.

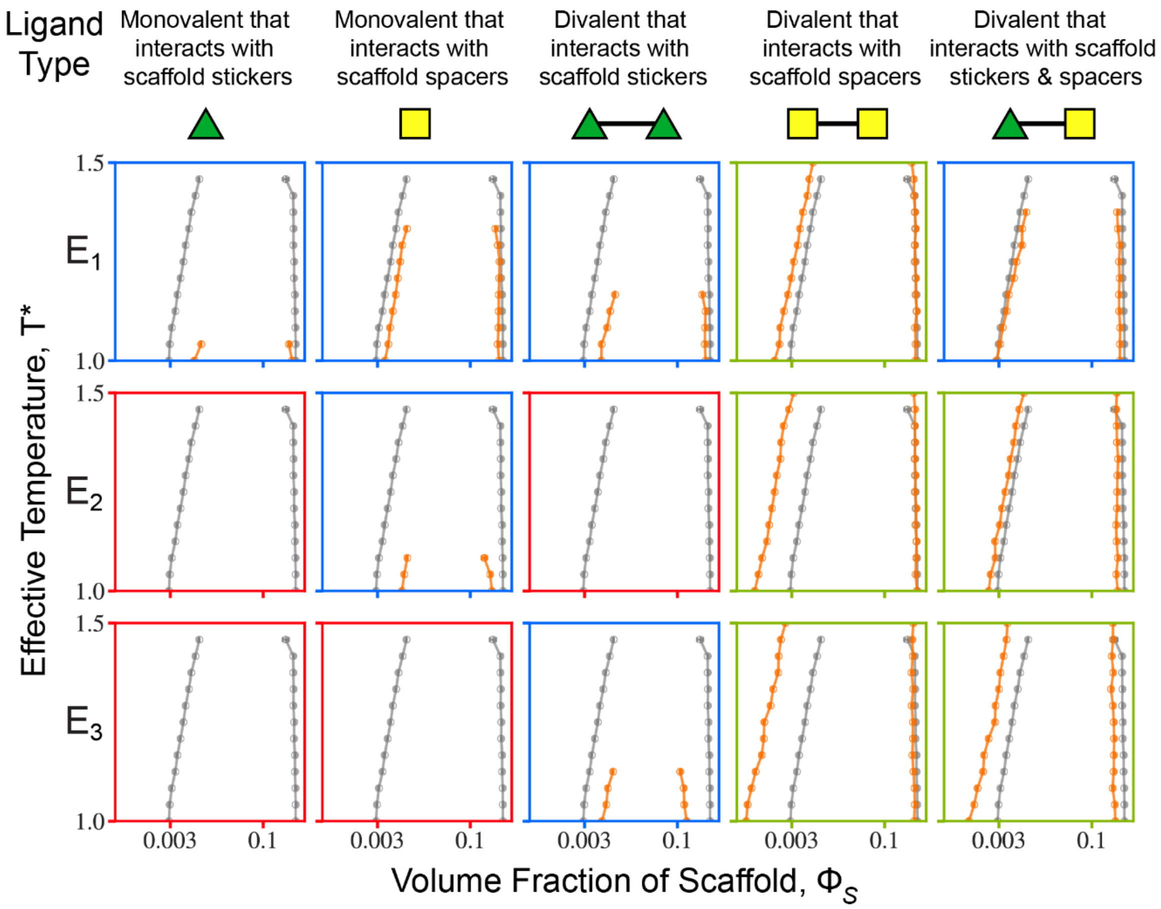

**Fig. S5.** Changes to $c_A^L$ as a function of ligand concentration at $T^*\approx1.08$ for the divalent ligand that binds both stickers and spacers on the scaffold for different energy scales. The legend denotes the interaction strength between the ligand site that interacts with scaffold stickers and the interaction strength between the ligand site that interacts with scaffold spacers. Shaded regions represent whether the ligand destabilizes (red), does not change (green) or promotes (blue) scaffold phase separation. Data are plotted only if the system undergoes phase separation, i.e., the width of the two-phase regime satisfies the criterion $\left(c_B^L - c_A^L\right) > 0.15$.

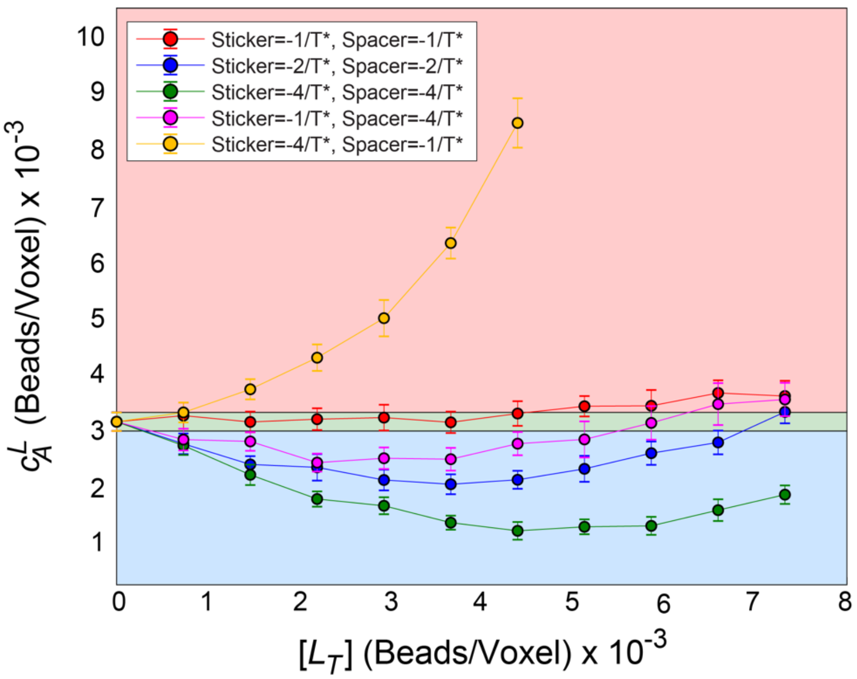

**Table S1.** Scaffold mutations associated with disease whose phase behavior changes from that of WT

| Scaffold | Mutation | Disease | Effect | Citation |
|---|---|---|---|---|
| **FUS** | G156E, R244C | Amyotrophic lateral sclerosis | Accelerates liquid-to-solid transition of FUS droplet | Patel et al. *Cell* 2015 (3), Murakami et al. *Neuron* 2015 (4) |
| **HnRNPA1** | D262V | Amyotrophic lateral sclerosis | Accelerates liquid-to-solid transition of hnRNPA1 droplet | Molliex et al. *Cell* 2015 (5) |
| **Huntingtin** | Polyglutamine expansion | Huntington's disease | Promotes phase separation<br><br>Promotes liquid-to-solid transition | Crick et al. *PNAS* 2013 (6)<br><br>Peskett et al. *Mol Cell* 2018 (7) |
| **SPOP** | W131G, F133V | Prostate cancer | Destabilizes phase separation | Bouchard et al. *Mol Cell* 2018 (8) |
| **Tau** | tau441 P301L, tau441 P301S, tau441 ΔK280, tau441 A152T | Alzheimer's disease | Makes droplets under conditions in which WT tau441 does not | Wegmann et al. *EMBO J* 2018 (9) |
| **TDP-43** | A321G, Q331K, M337V, A326P, M337P | Amyotrophic lateral sclerosis | Destabilizes phase separation by disrupting self-interactions | Conicella et al. *Structure* 2016 (10) |
| **UBQLN2** | P506S, P506T, P506A, T487I, P497L, P497H, P497S | Amyotrophic lateral sclerosis | Promotes phase separation and destabilizes dissolution | Dao et al. *Structure* 2019 (11) |

**Table S2.** Known ligands and their effect on scaffold phase behavior

| Scaffold | Ligand | Effect | Citation |
|---|---|---|---|
| **FUS** | Kapβ2 | Abolishes FUS liquid-liquid phase separation and disaggregates FUS fibrils by strongly binding PY-NLS and weakly binding FUS stickers | Guo et al. *Cell* 2018 (12), Yoshizawa et al. *Cell* 2018 (13) |
| **FUS R244C, FUS G156E *ALS related mutations** | Kapβ2 | Recovers WT FUS phase separation properties when disease related mutants promote phase separation (R244C) and aging (G156E) | Niaki et al. *Mol Cell* 2020 (14) |
| **FUS G156E FUS P525L, FUS R521C *ALS related mutations** | Lipoamide Lipoic acid | Delayed fibril formation and condensate hardening that is accelerated with the G156E mutation in vitro<br><br>Reduced motor defects in flies expressing P525L or R521C FUS | Wheeler et al. bioRxiv 2019 (15) |
| **FUS** | PAR polymer | PAR is necessary for FUS recruitment at DNA damage sites in vivo<br><br>Promotes FUS phase separation in vitro | Patel et al. *Cell* 2015 (3), Altmeyer et al. *Nat Comm* 2015 (16)<br>Patel et al. *Cell* 2015 (3) |
| **FUS** | $PR_{30}$ | Promotes FUS phase separation in vitro and enhances liquid-to-solid transition | Boeynaems et al. *Mol Cell* 2017 (17) |
| **FUS prion-like domain** | $[RGRGG]_5$ | Promotes FUS prion-like domain phase separation in vitro | Kaur et al. bioRxiv 2020 (18) |
| **G3BP1/RNA** | Caprin-1 | Enhances stress granule formation in cells<br><br>Promotes G3BP1 phase separation in vitro | Kedersha et al. *JCB* 2016 (19), Yang et al. *Cell* 2020 (20)<br><br>Guillén-Boixet et al. *Cell* 2020 (21), Yang et al. *Cell* 2020 (20) |
| **G3BP1/RNA** | TIA1 | Enhances stress granule formation in cells<br><br>Promotes G3BP1 phase separation in vitro | Yang et al. *Cell* 2020 (20)<br><br>Yang et al. *Cell* 2020 (20) |
| **G3BP1/RNA** | USP10 | Destabilizes stress granule formation in cells | Kedersha et al. *JCB* 2016 (19), Sanders et al. *Cell* 2020 (22) |
| **HnRNPA1 ∆hexa** | BSA | Destabilizes phase separation and reduces concentration of hnRNPA1 ∆hexa in the dense phase with increasing BSA concentration | Protter et al. *Cell Reports* 2018 (23) |

| | | | |
|---|---|---|---|
| **Huntingtin** | Profilin | Reduces huntingtin aggregation in cells<br><br>Destabilizes huntingtin phase behavior in vitro | Shao et al. *MCB* 2008 (24)<br><br>Posey et al. *JBC* 2018 (25) |
| **Huntingtin** | SH3GL3 | Promotes huntingtin aggregation in cells | Sittler et al. *Mol Cell* 1998 (26) |
| **PGL-3** | mRNA | Promotes PGL-3 phase separation such that it phase separates at its physiological concentration | Saha et al. *Cell* 2016 (27) |
| **$SH3_5/PRM_5$** | Heparin | Promotes phase separation but then destabilizes phase separation at high heparin concentrations | Ghosh et al. *PNAS* 2019 (28) |
| **$SH3_5/PRM_5$** | Lysozyme | Destabilizes phase separation | Ghosh et al. *PNAS* 2019 (28) |
| **TDP-43** | PAR polymer | Promotes TDP-43 liquid-liquid phase separation in vitro | McGurk et al. *Mol Cell* 2018 (29) |
| **UBQLN2** | Ub | Destabilizes UBQNL2 liquid-liquid phase separation | Dao et al. *Mol Cell* 2018 (11) |
